# Leveraging roadless areas to close strict protection gaps in the EU

**DOI:** 10.1101/2025.07.15.664891

**Authors:** Riccardo Testolin, Georg Hähn, Michele Di Musciano, Francesco Maria Sabatini, Alessandro Chiarucci

## Abstract

Strictly protected areas are critical to biodiversity conservation, yet only 4% of the EU’s land currently meets this designation, falling short of the 10% target outlined in the EU Biodiversity Strategy for 2030. Roadless areas, i.e., portions of land 1 km farther from linear infrastructure, are increasingly recognized for their ecological value and potential role in closing this gap. Using OpenStreetMap data and environmental gradients derived from climate, soil, and topographic layers, we identified the extent of EU roadless areas and assessed their conservation value based on their capacity to complement the existing strict protection network in terms of environmental representativeness. We found that 11.5% of EU land is roadless, with only 18.9% of this already under strict protection. Integrating all roadless areas would raise strict protection coverage to 12.7% and improve representation of abiotic environmental space from 78.8% to 94.4%. However, most roadless areas are extremely fragmented and redundant with respect to their abiotic features. While roadless areas offer a critical opportunity to expand the strict protection network, they cannot solely meet conservation goals. A balanced strategy that includes restoration, coordination across governance levels, and prioritization based on other biodiversity features is essential to effectively meet EU conservation commitments.

## Introduction

Area-based conservation is the most widely advocated tool to combat the current biodiversity crisis (Maxwell et al., 2020; Watson et al., 2014). Over the last decades, several global initiatives have emerged to strengthen this strategy, culminating in the adoption of the Kunming-Montreal Global Biodiversity Framework. This framework defines long-term global biodiversity goals for 2050 alongside short-term targets for 2030, among which is the protection of 30% of the planet’s surface (Convention on Biological Diversity, 2022). The European Union (EU) incorporated this target in its Biodiversity Strategy for 2030, which includes provisions to protect 30% of the EU’s ecosystems (European Commission, 2020). Additionally, the strategy mandates that at least 10% of each member state’s surface should be placed under strict protection. This framework has been reinforced by the EU Nature Restoration Law, which mandates legally binding restoration targets for degraded ecosystems while emphasizing the importance of achieving area-based conservation objectives (European Commission, 2022). However, the EU is currently falling short of its own commitments, with protected areas covering only 26% of land (European Environment Agency, 2024), with less than 4% under strict protection (Cazzolla Gatti et al., 2023). The gap in strictly protected areas is particularly critical, as these are essential for preserving wilderness (Dudley, 2008), defined as areas governed by natural processes that are “[…] *unmodified or only slightly modified and without intrusive or extractive human activity, settlements, infrastructure, or visual disturbance*” (European Commission, 2013). This is a fundamental target for biodiversity conservation, given the catastrophic decline in wilderness across the world’s biomes (Potapov et al., 2017; Watson et al., 2016).

While not necessarily corresponding to true wilderness, roadless areas – defined as natural and semi-natural regions largely devoid of roads (Selva et al., 2011) – represent some of the EU’s most undisturbed landscapes (Kati et al., 2023). Transport infrastructure is a major threat to biodiversity and ecosystem functioning (Torres et al., 2016). Roads extend anthropogenic disturbance into natural environments by increasing habitat fragmentation, noise, pollution, as well as the risk of fire, biological invasions, road kills and behavioural changes in many taxa (Laurance et al., 2014; Selva et al., 2011; Trombulak & Frissell, 2000). By granting access to remote areas, roads also promote “contagious development”, allowing commercial forestry and triggering habitat degradation and loss (Ibisch et al., 2016). Due to their relatively high conservation value and reduced societal conflict risks stemming from limited accessibility (Psaralexi et al., 2017), roadless areas are emerging as important candidates for the establishment of new strictly protected areas (Brackhane et al., 2019; Ibisch et al., 2016; Kati et al., 2023).

While the protection of roadless areas has been advocated by many studies at global (Ibisch et al., 2016), continental (Psaralexi et al., 2017), and national scale (Brackhane et al., 2019; Kati et al., 2023), there is limited guidance on which areas should be prioritized. Expanding protected areas should not be limited to designating additional land but should rather address deficiencies in existing networks.

The planning of the current system of protected areas has largely been opportunistic and politically driven, with many sites being established based on compromise solutions meant to minimize social conflicts without a systematic planning. Protected areas are disproportionately located at high-elevation and in infertile areas (Joppa & Pfaff, 2009), a bias that is particularly true for strictly protected areas (Cazzolla Gatti et al., 2023). As a result, many species and ecosystems are underrepresented in the current protected area network (Hoffmann et al. 2018; Hanson et al., 2020; Sabatini et al., 2020; Mouillot et al., 2024). Expanding strictly protected areas should therefore prioritize currently overlooked biodiversity elements to enhance representativeness while aiming to the 10% area target.

The prioritization of sites should also account for the multiple scales at which conservation targets are defined and implemented (Hoffmann et al., 2018). As a matter of fact, international conservation strategies are enacted by national or subnational governments by integrating global and continental priorities with local needs and ensuring a fair distribution of conservation efforts (Secretariat of the Convention on Biological Diversity, 2004; Stark et al., 2022). However, independent national and local actions aimed at achieving area targets can result in inefficiencies and redundancies. For example, a country may overlook the protection of a habitat that is well represented within its own borders but rare and insufficiently protected at the continental level. Conversely, countries may prioritize protecting a given habitat because it is locally rare, even if it is already well covered at the continental scale or, more in general, they may focus on areas of low international importance. Historically, the lack of cross-country coordination has hindered progress toward regional conservation goals (Brooks et al., 2004; Montesino Pouzols et al., 2014), reducing the overall efficiency of protected areas for biodiversity conservation (Hermoso et al., 2020; Moilanen et al., 2013). Effective solutions to achieve conservation targets should incorporate both national and international perspectives (Hermoso, 2015; Hermoso et al., 2020; Moilanen et al., 2013).

In systematic conservation planning, protected area expansion is traditionally guided by the presence of endangered species or habitats, with the aim to cover a given portion of their range (Margules & Pressey, 2000). However, this approach is sensitive to the selected biodiversity features (Wilson et al., 2005) and may fail to ensure long-term persistence under projected climate change due to loss of climatic suitability (Araújo et al., 2011). Moreover, significant knowledge gaps persist regarding the distribution of species of high conservation value – the so-called “Wallacean shortfall” (Hortal et al., 2015) – even within areas where biodiversity is relatively well-known, such as the EU (Marshall et al., 2024; Navarro et al., 2024). Species requiring the greatest protection are often the ones we know the least about, complicating distribution modelling (Jeliazkov et al., 2022; Lomba et al., 2010).

Consequently, relying solely on known biodiversity features in systematic conservation planning may overlook areas with significant potential conservation value, such as undisturbed habitats hosting specific and likely understudied biotas. Growing research advocates shifting the focus to abiotic environmental space within open-ended management frameworks (Bergin et al., 2024; Brunbjerg et al., 2017). Indeed, environmental gradients underpin ecological processes and drive broad-scale biodiversity patterns (Moura et al., 2016). Thus, emphasizing abiotic envelope representation in a process-oriented perspective when planning protection network expansions can address these limitations and help maintain biodiversity over time (Bergin et al., 2024). The potential contribution of regions with lower human disturbance like roadless areas to enlarge the abiotic envelope represented in existing protected area networks in Europe is still unknown.

Here, we leverage open-source, high-resolution infrastructure data from OpenStreetMap (OSM; OpenStreetMap contributors, 2024) to explore the potential contribution of European roadless areas to achieving strict protection targets. Specifically, we addressed three research gaps by 1) estimating the amount, distribution, and current inclusion of EU roadless areas in the existing network of strictly protected areas; 2) assessing how well the current strict protection network and roadless areas are representative of the variety of abiotic environmental gradients occurring in the EU; 3) identifying the conservation value of roadless areas based on their potential to complement the existing strict protection network in representing abiotic environmental gradients, both at the national and EU level.

## Methods

### Identification of roadless and wilderness areas

Roadless areas in the EU were identified using data from OSM (OpenStreetMap contributors, 2024), accessed via the *osmdata* package (Padgham et al., 2017) in R v4.4.1 (R Core Team, 2022). While OSM data is entirely user-contributed, the EU network has been shown to be virtually complete, and its quality is comparable to other official or proprietary datasets (Barrington-Leigh & Millard-Ball, 2017). Roadless areas were defined as regions more than 1 km away from paved roads and railways, being 1 km an average threshold beyond which areas can be considered significantly unaffected by linear transport infrastructure (Ibisch et al., 2016).

To delineate roadless areas, 1-km spatial buffers were created around each linear feature while excluding tunnels and bridges, based in OSM tags (Ramm, 2022; Table S1). All terrestrial patches not intersected by the buffers were consequently extracted. To avoid including polygons with high perimeter-to-area ratios (i.e., narrow or edge-like features) and to increase compactness, we applied a 300-m inward spatial buffer followed by a 300-m outward one (Brackhane et al., 2019) (Figure S1).

Additionally, to simplify the dataset and exclude scattered fragments, we removed the polygons smaller than 1 km². Remaining roadless polygons falling within inland water bodies or containing more than 10% cropland or built-up area were excluded. This filtering was based on 10-m resolution land cover data for 2021 (Zanaga et al., 2022), accessed via Google Earth Engine (Gorelick et al., 2017) using the *rgee* package (Aybar, 2023).

While roadless areas as defined above are largely inaccessible, they are not entirely free from human influence. To identify a subset of roadless areas with minimal human presence (‘wilderness areas’, hereafter), we repeated the workflow, this time including all linear features such as unpaved roads, suspended infrastructures (e.g., cable cars), artificial waterways, power lines, and pipelines (Table S1). The extent and distribution of roadless and wilderness areas were analysed across countries and size classes (>1 km^2^, >5 km^2^, >10 km^2^, >50 km^2^, >100 km^2^, >1000 km^2^) following (Ibisch et al., 2016).

### Identification of strictly protected areas

We utilized data from the World Database of Protected Areas (WDPA), a joint initiative of the UN Environment Programme (UNEP) and the International Union for Conservation of Nature (IUCN), which represents the most comprehensive global database of terrestrial and marine protected areas (UNEP-WCMC & IUCN, 2024). WDPA data for the EU – which include nationally designated areas found in the European Common Database on Designated Areas (CDDA), as well as international designation and other effective area-based conservation measures – were downloaded using the *wdpar* package (Hanson, 2022). The raw dataset was refined by excluding marine protected areas and resolving polygon overlaps.

Marine protected areas were removed using information from the dataset and performing a manual assessment. First, we excluded areas marked as marine in the respective attribute column or classified under marine designations (e.g., Protected Marine Area, National Marine Park). Next, we cleaned the geometries using the *wdpa_clean* function, which resolves overlapping polygons by retaining those associated with more effective management categories, as per the IUCN classification of protected areas (Dudley, 2008). Since Croatia’s data lacked IUCN category information, we classified its polygons following (Underwood et al. (2014) before applying this cleaning step. The polygons were then cut to the countries’ extents. Finally, the cleaned dataset was manually inspected in QGIS (QGIS Development Team, 2024) to identify and remove any remaining marine protected areas based on their location and name. Terrestrial strict protected areas were defined as those belonging to the following IUCN protected area categories: Ia – strict nature reserve, Ib – wilderness area, and II – national park. These polygons were subsequently used to calculate the proportion of strictly protected roadless and wilderness areas.

### Definition of the environmental space

To characterize the main environmental gradients underpinning ecological processes and biodiversity patterns, we collated a parsimonious set of abiotic (i.e., climatic, edaphic, and topographic) factors relevant to species survival and growth. Although we did not explicitly target any specific species group when selecting the variables, our set primarily reflects those influencing plant life cycle, as plants are key determinants of terrestrial ecosystem structure and functioning (Freschet et al., 2021; Schuldt et al., 2018; Cervellini et al. 2021). All variables were derived from global geospatial raster data layersat a 30 arc-second resolution (i.e. north-south resolution of ∼1 km and east-west resolution of ∼0.7 km).

For climate, we selected monthly and seasonal extremes in temperature and precipitation, namely the maximum temperature of the warmest month (bio5), minimum temperature of the coldest month (bio6), precipitation of the warmest quarter (bio18), and precipitation of the coldest quarter (bio19) (Mckenney et al., 2007; Zellweger et al., 2016). Climatic extremes are tightly linked to species demography and distribution and are widely employed in predictive models (Stewart et al., 2021; Zimmermann et al., 2009). We also included site water balance (swb) and snow cover days (scd) (Zellweger et al., 2016). The former estimates water availability, while the latter represents the number of days per year the ground is covered with snow (Brun et al., 2022). Additionally, we selected downwelling solar radiation (rsds), which estimates solar energy availability and influences local microclimates and vegetation patterns in high-latitude and high-elevation environments (Brun et al., 2022; Sabatini et al., 2021). We also included potential net primary productivity (npp), given its well-established relationship with biodiversity (van Ruijven & Berendse, 2005). All these climatic predictors were retrieved from CHELSA v2.1 (Karger et al., 2017).

For edaphic variables, we included soil pH, soil organic carbon (soc), and the percentages of silt and clay at a depth of 5–15 cm, derived from SoilGrids v2.0 (Poggio et al., 2021). Topsoil factors strongly influence species distribution and richness patterns (Delgado-Baquerizo et al., 2020; Spohn et al., 2023; Zellweger et al., 2016). Finally, we used terrain roughness from Amatulli et al. (2018) as a measure of local topographic heterogeneity and microhabitat availability. All variables were standardized to 0 mean and 1 standard deviation.

We removed pixels lacking edaphic variable data (i.e., urban areas, inland waters, glaciers, and bare surfaces) and conducted a principal component analysis on the standardized variables. The loadings of the individual variables were used to identify the main axes of variation. The first two principal components (PCs) together explained 67% of the variance in the environmental data and were selected to represent the abiotic environmental space of terrestrial habitats (‘environmental space’, hereafter) (Figure 1). The first axis (PC1), which explained 51% of the variance, primarily represented a gradient of air temperature and soil chemical properties. The second axis (PC2), explaining 16% of the variance, was largely associated with precipitation, terrain roughness, and net primary productivity (Figure 1 b,c).

**Figure 1.**
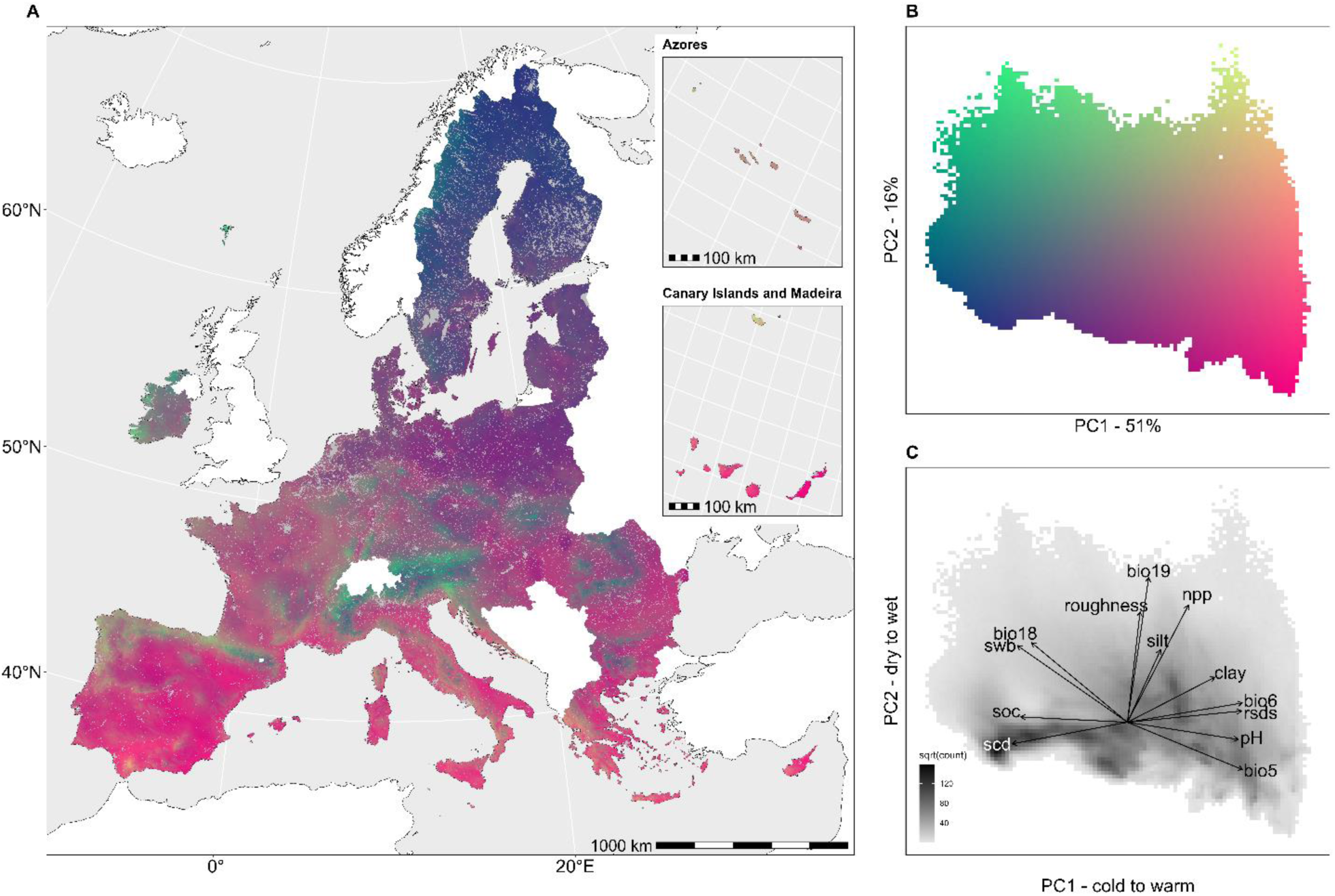
Environmental space of the EU. A) EU territory coloured according to its position in a two-dimensional environmental space defined by the first two axes of a principal component analysis (PCA) of 13 scaled environmental variables. An interactive version of this map is available at: https://ee-riccardotestolin-367410.projects.earthengine.app/view/roadless-areas-in-the-european-union. B) the PCA space is divided into a 100 × 100 grid; each cell is coloured according to its position in the space (PC1 and PC2). C) Density of the environmental space shown as cells shaded by the square root of the number of pixels (i.e. geographic locations) falling within that cell. Variable loadings on the first two PCA axes are shown, where arrow direction and length represent the direction and strength of correlation, respectively.

To evaluate the distribution of EU territory within the environmental space, we subdivided the principal component plane into a uniform 100 × 100 grid and counted the number of 30 arc-second resolution pixels from the EU geographical space falling within each cell. Each grid cell represents a distinct portion of the environmental space, which has a corresponding geographic extent given by the sum of the pixels’ areas. Finally, PC1 and PC2 values were mapped onto the EU geographical space to spatialize the distribution of pixels in the environmental space and help with its interpretation. The top-left portion of the environmental space corresponds to mountain areas, while the top-right reflects areas with stronger Atlantic influence. The centre-bottom portions of the environmental space represent abiotic conditions that cover wide extents of the EU territory, corresponding to high-altitude (bottom-left) and continental (centre-right) regions. The bottom-right portion corresponds to Mediterranean regions (Figure 1 a,b).

Variable abbreviations: bio5 = maximum temperature of the warmest month, bio6 = minimum temperature of the coldest month, bio18 = precipitation of the warmest quarter, bio19 = precipitation of the coldest quarter, npp = net primary productivity, rsds = downwelling shortwave radiation, scd = snow cover days, soc = soil organic carbon, swb = soil water balance.

### Analysis of environmental representativeness

To assess how the inclusion of wilderness and roadless areas would increase the environmental representativeness of the strict protection network we considered three configurations: 1) the current network of strictly protected areas only, and two hypothetical networks also incorporating 2) all wilderness areas and 3) all roadless areas. Notably, we assessed the increase in representativeness incrementally, by first expanding the strict protection network with all wilderness areas – which are fewer and virtually devoid of human presence – and then with all remaining roadless areas. To achieve this, we clipped the environmental geospatial raster data layers using the polygons corresponding to each configuration and projected the resulting values onto the environmental space. For each grid cell in the environmental space, we calculated the proportion of pixels covered by each configuration, distinguishing grid cells where 1) at least 10% of the pixels, 2) less than 10% of the pixels, and 3) no pixels were covered. The 10% threshold was selected to define adequate coverage within the environmental space, aligning with the 10% area target set by the EU Biodiversity Strategy for 2030. These analyses were repeated for each country individually by subsetting the EU environmental space to their respective extents.

### Conservation value

Based on the environmental representativeness, we determined the conservation value of wilderness and roadless areas that are currently falling outside the strict protection network. First, we projected the proportion of pixels in each grid cell of the environmental space covered by the current network of strictly protected areas – thus ranging from 0% to 100% – onto the geographical space of wilderness and roadless areas outside the strict protection network, based on the spatialized PC1 and PC2 scores (Figure 1). This provided a measure of protection priority for each pixel in the geographical space, with low percentages indicating portions of the environmental space encompassed by wilderness and roadless areas that are currently underrepresented by the strict protection network and thus have higher priority. Next, we derived a measure of redundancy for wilderness and roadless areas outside the strict protection network. We did this by calculating, for each pixel in the geographical space, the absolute percentage change by which the coverage of the corresponding grid cell in the environmental space would increase after including all wilderness and roadless polygons in the strict protection network. This measure – which is necessarily positive due to the additive contribution of wilderness and roadless areas to the representation of the environmental space – ranges from >0% to 100%, where high values indicate portions of the environmental space that are overrepresented in wilderness and roadless areas, and that are this redundant. We then summarized the priority and redundancy scores by calculating their mean for each wilderness and roadless polygon. A 10% threshold was again used to distinguish between high and low priority and redundancy, allowing classification of wilderness and roadless polygons into three conservation value categories at the EU level: 1) low priority, 2) high priority and high redundancy, and 3) high priority and low redundancy. Low priority polygons – regardless of redundancy – occupy portions of the EU environmental space already adequately represented by the current strict protection network (i.e., ≥10% of pixels within each grid cell). In contrast, high priority and high redundancy polygons are areas in environmental conditions that are underrepresented in the strict protection network, but that would become overrepresented if all respective wilderness or roadless polygons were included in the network. Finally, high priority and low redundancy polygons identify areas currently underrepresented in the EU environmental space that would remain so even after inclusion in the strict protection network, thus representing the class with the highest conservation value. The same analyses were repeated for each EU country to determine the conservation value at the national level.

Finally, we synthesized the abovementioned classification into a more actionable framework to assess the contribution of an environmentally balanced selection of wilderness and roadless areas toward achieving the 10% area target for strict conservation. To this end, we further grouped these areas based on their EU and national conservation values into three classes: 1) core value areas (high priority and low redundancy at both national and EU levels), which should be included in protected area expansion; 2) conditional value areas (inconsistent combinations, e.g., high priority and low redundancy at only one level), whose inclusion should be evaluated on a case-by-case basis; and 3) low value areas (low priority at both levels), which can be deprioritized for strict protection. For each conservation value class at both the national and EU levels, we calculated the proportion of wilderness and roadless areas in that class relative to the total area of wilderness and roadless land falling outside the strict protection network, as well as the potential percentage increase in strictly protected area coverage if those areas were included.

## Results

Roadless areas in the EU cover 476,583 km², corresponding to 11.5% of the land surface area of EU member states. Wilderness areas, totalling 193,745 km², only account for 4.7% of EU surface and represent 40.7% of the roadless extent. More than three-quarters (76.6%) of the EU’s roadless areas are concentrated in Sweden (30.8%), Finland (17.3%), Spain (14.5%), Romania (9.3%), and Italy (4.8%). These same countries encompass 84.5% of the EU’s wilderness areas, most of which are in Sweden (43.6%) and Finland (21.7%). Belgium has no wilderness areas but retains small roadless areas (0.01%), while Luxembourg and Malta have none (Figure 2a, b; Table S2).

**Figure 2.**
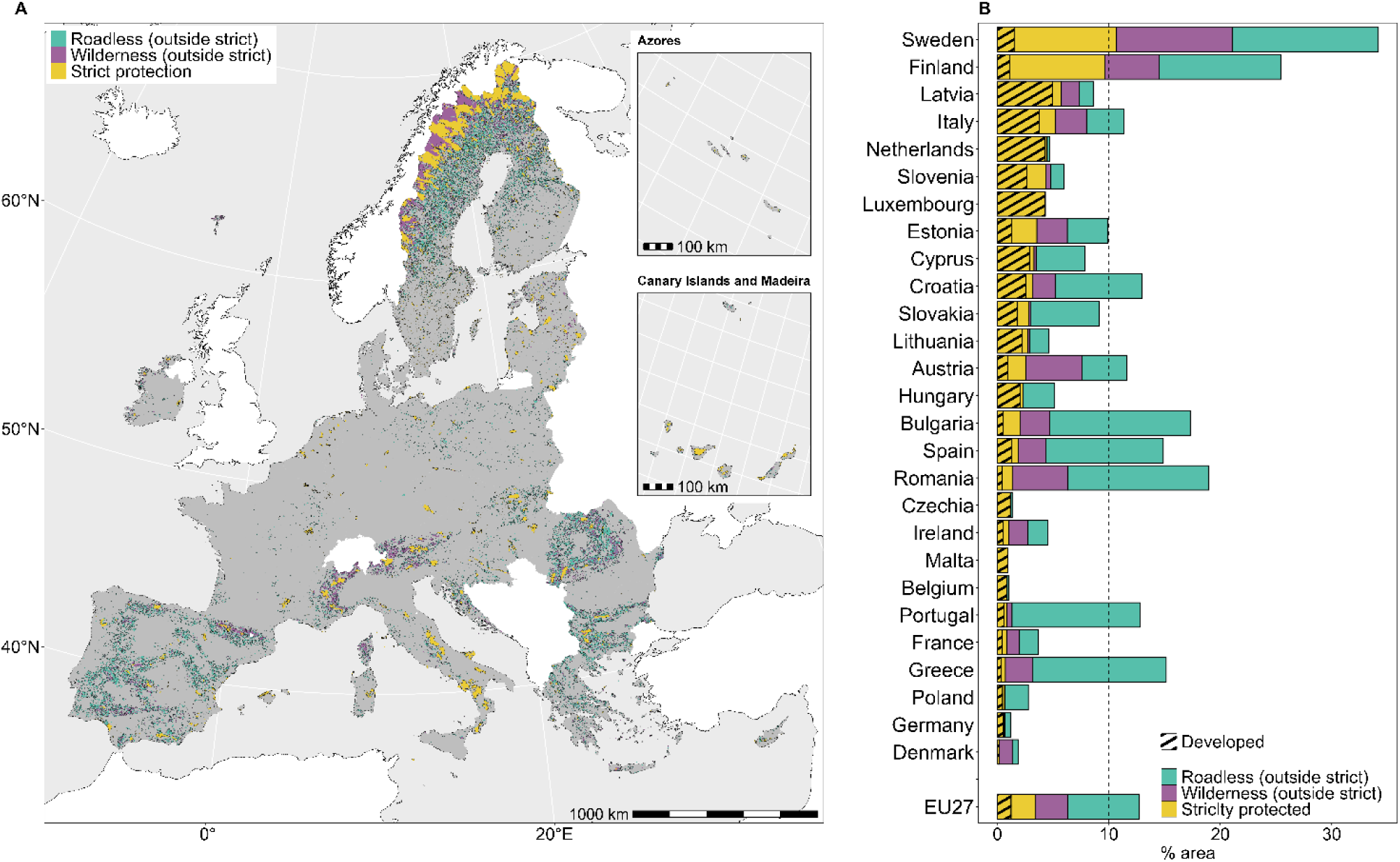
Distribution and proportion of roadless, wilderness, and strictly protected areas in the EU. A) spatial distribution of roadless, wilderness and strictly protected areas (IUCN categories I & II) across the 27 EU countries. An interactive version of this map is available at: https://ee-riccardotestolin-367410.projects.earthengine.app/view/roadless-areas-in-the-european-union. B) proportion of strictly protected areas and additional roadless and wilderness areas currently outside the strict protection network for each country and for the EU. The dashed line represents the 10% area target for strict protection. The stripes represent the developed fractions of strictly protected areas, i.e., those that were not identified as wilderness or roadless.

**Figure 3.**
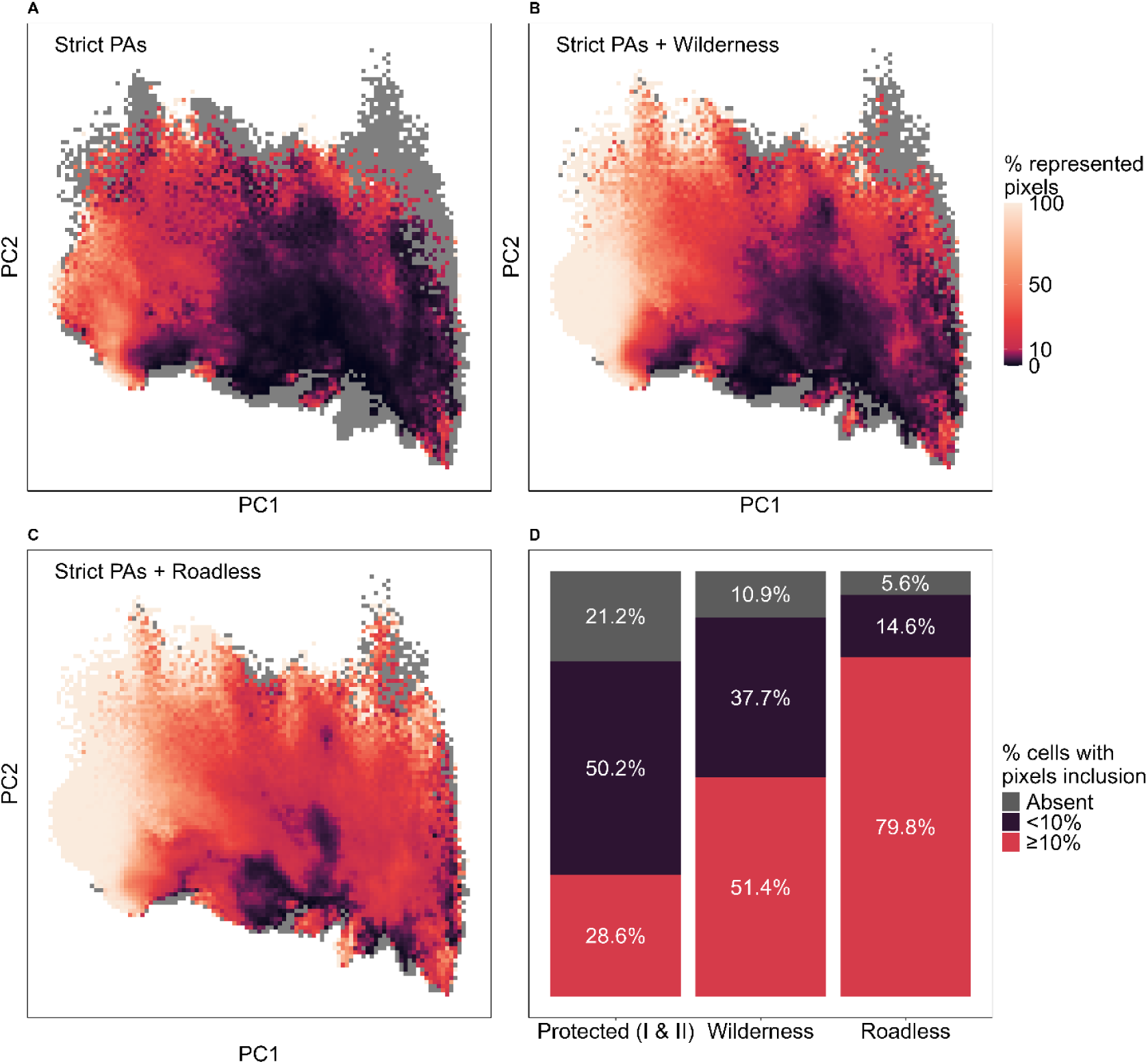
Coverage of the EU environmental space by strictly protected, wilderness and roadless areas. Proportion of EU territory (% 30-arc second resolution pixels from the EU geographical space) within each grid cell of the environmental space represented by different configurations of the network of strictly protected areas: A) strictly protected areas (IUCN categories I & II), B) strictly protected and all wilderness areas, and C) strictly protected and all roadless areas. D) proportion of the EU environmental space (% of grid cells in the environmental space) covered by the current strict protection network and the hypothetical configurations including all wilderness and roadless areas, respectively. A cell was considered overrepresented in a given configuration when the proportion of pixels included was above the 10% threshold.

Roadless and wilderness areas are fragmented into 22,701 and 9,967 patches, respectively. The majority (55.6%) of roadless fragments are smaller than 5 km², while 27.8% are larger than 10 km², and only 3% exceed 100 km². Just 31 (0.1%) areas are larger than 1,000 km², while making up 23% of the total roadless area in the EU. Similarly, 63.3% of wilderness fragments are smaller than 5 km², 19.9% are larger than 10 km², and only 2% exceed 100 km², with just 22 (0.2%) surpassing 1,000 km² and making up 39% of the total wilderness area. Fragment sizes vary considerably across countries, with the largest patches primarily found in Finland, Sweden, Romania, and Spain (Table S3).

Regarding current protection, 90,126 km² (18.9%) of EU roadless areas and 74,233 km² (38.3%) of wilderness areas are already included in the strict protection network. The countries with the highest proportion of roadless areas within the strict protection network are Slovenia (51.8%), the Netherlands (35.6%), and Finland (35.2%), whereas those with the lowest proportions are Greece (2.5%), and Portugal (2.2%). Notably, no roadless area is included in the strict protection network of Belgium. As for wilderness areas, the highest proportions within the strict protection network are found in Hungary (82.9%), Slovakia (81.2%), and Slovenia (77.3%), while the lowest are in Romania (7.8%), Greece (7.2%), and Denmark (1%).

If all wilderness areas were placed under strict protection, the total strictly protected land in the EU would rise from 3.4% to 6.3%, and from 3.4% to 12.7% if all roadless areas were included. Nine countries have enough roadless area left to expand the current strict protection network and exceed the 10% land protection target, namely Finland (25.4%), Romania (18.9%), Bulgaria (17.3%), Greece (15.2%), Spain (14.9%), Croatia (13%), Portugal (12.8%), Austria (11.6%), and Italy (11.3%). Sweden, with 10.5% of its land already under strict protection, has already reached the target. Conversely, other countries – particularly Denmark, Germany, Poland, France, Belgium, Malta, and Czechia – are still far from the 10% target and have little or no remaining roadless areas available (Figure 2b, Table S4).

The current network of strictly protected areas encompasses 78.8% of the EU environmental space. Within this fraction, 28.6% of the total environmental space - mainly corresponding to high-elevation and high-latitude regions – is adequately or overrepresented, passing the 10% threshold, while 50.2% is underrepresented, being covered for less than 10%. Integrating all wilderness areas into the network would increase overall coverage of the environmental space to 89.1%, with 51.4% of the total environmental space adequately or overrepresented and 37.7% underrepresented. Further inclusion of all roadless areas would leave only 5.6% of the EU environmental space not represented and 14.6% underrepresented, while 79.8% would be adequately or overrepresented. In both cases, the inclusion of wilderness and roadless areas would disproportionately favour high-elevation and high-latitude regions (Figure 1; 3).

Across individual countries, the current strict protection network covers between 0.4% (Malta) and 84.3% (Sweden) of the environmental space. In some countries, the inclusion of wilderness areas would result in minimal coverage increases, such as +0.2% in the Netherlands, +0.3% in Czechia, and +0.4% in Hungary, whereas in others, the increase would be substantial, e.g., +46% in Denmark, +41.5% in Greece, and +31.6% in Bulgaria. If all roadless areas were included, the smallest coverage increases would be slightly higher, e.g., +2.1% in the Netherlands, +2.2% in Belgium, and +5.8% in Sweden, while the largest increases would also be greater, e.g., +61.3% in Greece, +56.7% in Denmark, and +55.8% in Croatia (Figure S2).

Roadless (hereafter including wilderness) areas with high priority and low redundancy at both the EU and national levels were concentrated mainly in the Maritime Alps, the mountains of Corsica and Greece, as well as in the lowlands of France, Germany, Poland, Scandinavia and Baltic countries.

Thus, these areas were assigned to the core value group. Roadless areas of Mediterranean mountains (e.g. Apennines, Balkans) exhibited high priority and low redundancy at the EU level but low priority at the country level. Conversely, most areas of the Alps were classified as high priority and low redundancy at the country level but low priority at the EU level. Most roadless areas of Bulgaria, Portugal, Romania and Spain showed high priority at both country and EU level but also high redundancy. All the abovementioned areas were assigned to the conditional value group. Finally, a large fraction of roadless areas in northern Scandinavia, as well as in the Alps, exhibited low priority at both the EU and national levels and were thus classified as low value (Figure 4a, b).

**Figure 4.**
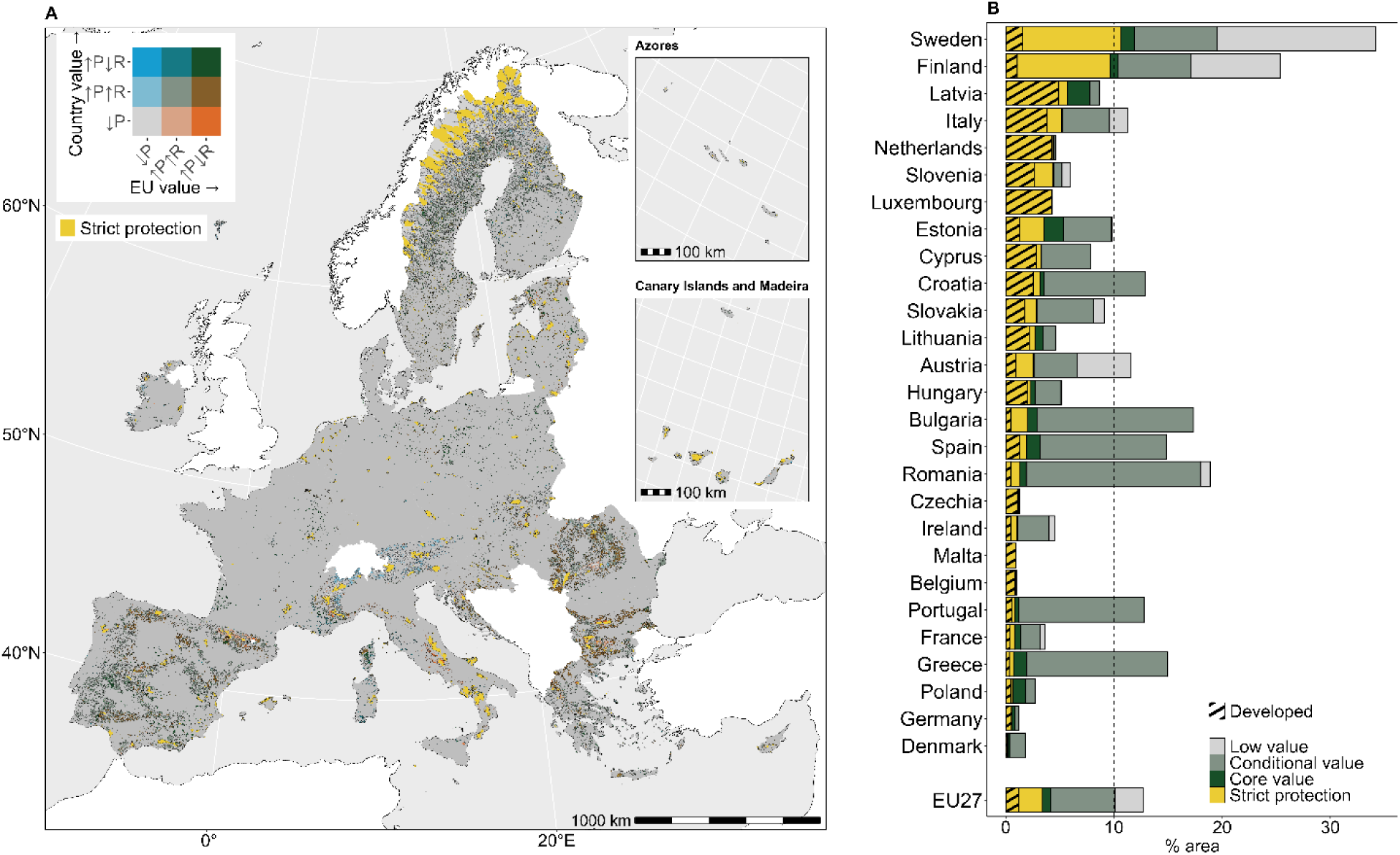
Conservation value of roadless (including wilderness) areas. A) classification of roadless areas based on mean priority and redundancy scores at the EU and country levels. Legend abbreviations: ↓P = low priority, ↑P↑R = high priority & high redundancy, ↑P↓R = high priority & low redundancy. An interactive version of this map is available at: https://ee-riccardotestolin-367410.projects.earthengine.app/view/roadless-areas-in-the-european-union. B) proportion of strictly protected areas and additional roadless areas currently outside the strict protection network for each country and for the EU, classified according to conservation value. The stripes represent the developed fractions of strictly protected areas, i.e., those that were not identified as wilderness or roadless.

In the EU, only 7.9% of unprotected roadless areas together were classified as core value, while 28.3% were classified as low value. The remaining (63.2%) were classified as conditional value. Including all core and conditional value areas into the strict protection network would increase its cover to 10.1% of the EU land surface. The countries with the highest proportion of unprotected roadless areas of high conservation value were Latvia (70.4%), Poland (57.5%), and Germany (44.7%), while those with the highest proportion classified as low conservation value were Sweden (62.1%), Austria (55.3%), and Finland (52.3%). Italy and Austria, while possessing enough roadless area, would not meet the 10% target if low value areas were excluded (Figure 4b, Table S5).

## Discussion

Meeting the objective of strictly protecting 10% of EU’s land by 2030, as mandated by the European Biodiversity Strategy, requires a substantial expansion of the existing network of strictly protected areas. Roadless areas, being relatively free of anthropogenic disturbance, may help reaching this target while minimizing societal conflict risks, as long as they are effective at increasing the representativeness of the strict protection network. In this study, we developed a reproducible workflow to identify roadless areas across the EU and evaluated their conservation value at both the national and EU levels based on their representativeness of the environmental space.

We found that 11.5% of the EU’s land surface can be considered roadless under a 1 km road-effect buffer. This estimate is significantly lower than previous assessments, such as the 30% reported by Psaralexi et al. (2017) for the former EU with 28 member states or the 42% by Ibisch et al. (2016) for the whole European continent. Yet, differences in methodological approaches limit straightforward comparisons. While Psaralexi et al. (2017) excluded many low-traffic road categories, Ibisch et al. (2016) included all road types in their analyses. Our study, based on a fully reproducible and adjustable approach, applied a conservative yet balanced definition of roadless areas, including all types of roads accessible by vehicles and excluding footpaths and unpaved tracks. Furthermore, the continuous improvements in the completeness of OSM data (Novack et al., 2024), as well as infrastructure expansion in the EU over the last decade (Eurostat, 2024), may have contributed to lower the estimate of roadless area extent. In addition, more than half of the roadless and wilderness patches are smaller than 5 km², while only a minor fraction exceeds 100 km², mirroring previous findings (Ibisch et al., 2016; Psaralexi et al., 2017). This extreme fragmentation undermines the ecological integrity and conservation value of roadless aeras. Small and isolated fragments exhibit lower biodiversity, stronger edge effects and limited recolonization (Haddad et al., 2015; Ibisch et al., 2016). This emphasizes the need to integrate roadless areas into larger ecological networks and consider road removal, rewilding, or corridor establishment to enhance their functional connectivity (Jaeger et al., 2011; Perino et al., 2019; Selva et al., 2011).

The spatial distribution of roadless areas across countries is highly uneven. Northern and Eastern European countries, notably Sweden and Finland, retain the largest extents of roadless terrain, benefiting from vast forests and low population densities. Similarly, Romania and Bulgaria preserve substantial roadless areas, particularly in mountainous regions like the Carpathians and Rila– Rhodope massif (Selva et al., 2011). In the south, Greece and Spain exhibit relatively high shares of roadless area, located within the globally significant Mediterranean biodiversity hotspot (Myers et al., 2000). The distribution of roadless areas in Greece has been extensively investigated (Kati et al., 2020, 2023). This effort led to their integration into national legislation that designated several roadless mountain ranges as off-limits to infrastructure development, showing a direct application to conservation planning (Kati et al., 2022). In contrast, Western and Central European countries – including Germany, Czechia, and the Netherlands – have retained only minimal, scattered remnants, often comprising less than 1% of their land area, consistently with previous analyses (M. T. Hoffmann et al., 2024).

Less than half of roadless areas – corresponding to 4.7% of EU land surface – were classified as wilderness, defined here as regions devoid of any linear infrastructure, including forest roads, cable cars and power lines. By exhibiting relatively low levels of human presence and activities, remaining roadless and, especially, wilderness areas can be considered as “Anthropocene refugia”, i.e., landscapes that serve as strongholds for biodiversity and climate resilience in an era of pervasive change (Monsarrat et al., 2019). The role of these refugia in maintaining ecosystem processes, buffering climate change impacts, and supporting biodiversity in the long term, highlights the need for the establishment of effective protection initiatives in those sites. However, we found that only 18.9% of roadless and 38.3% of wilderness areas currently fall within the strict protection network. On the one hand, this means that a large fraction of the remaining roadless and wilderness areas are still exposed to potential infrastructure development. On the other hand, it also indicates considerable potential for expanding the network of strictly protected areas to encompass these regions.

Yet, the inclusion of roadless areas into the strict protection network should be evaluated carefully, rather than pursued indiscriminately, to avoid replicating longstanding spatial biases in protected areas. As with existing protected areas, roadless areas are often concentrated in remote, high-altitude regions (Joppa & Pfaff, 2009; Psaralexi et al., 2017), raising concerns that relying on roadless areas alone to meet area-based targets without accounting for environmental diversity could further reinforce these biases. Considering complementarity, i.e., the principle of selecting new conservation areas that add unique biodiversity features not yet represented, is paramount to effective expansion of protected area networks. Previous studies have shown that complementarity-based planning significantly enhances biodiversity representation while minimizing costs and land use conflicts (Di Marco et al., 2016; Rodrigues et al., 2004). Our results indicate that a strategic selection of roadless areas, guided by their representativeness in the environmental space, could aid the expansion of the EU strict protection network while incorporating underrepresented abiotic conditions. Based on environmental representativeness, we identified a subset of roadless areas that encompass environmental conditions that are seldom found within the current strict protection network.

Nonetheless, even their full inclusion would not ensure an adequate representation of the entire spectrum of the abiotic gradients we considered, which is why we assigned them core conservation value. These areas mainly include remnants of lowland ecosystems in Central Europe – which are widely recognized as underrepresented in current protected area networks (Cazzolla Gatti et al., 2023; Müller et al., 2020) and have limited residual extent to achieve prescribed area targets (Araújo & Alagador, 2024). Conversely, large roadless extents in the boreal and alpine zones are often characterised by environmental conditions which are already well-represented in the strict protection network. While these areas may still be worthy of strict protection for other ecological reasons beyond environmental representativeness – such as the presence of primary forests (Sabatini et al., 2020) or rare species (Dinerstein et al., 2024) – they were assigned low value in our framework and could therefore be put under less restrictive levels of protection (i.e., IUCN categories from III to VI), or employed as ecological buffers and corridors.

Most of the identified roadless areas (64%) were classified as conditional value and can be divided into two groups: 1) those that were assigned conflicting value rankings between the EU and national levels, and 2) those that fell in environmentally redundant zones, despite being underrepresented by the current strict protection network. The former highlights mismatches between national and international conservation targets. National conservation actions may prioritize locally valued features over broader regional priorities, potentially jeopardizing effectiveness (Montesino Pouzols et al., 2014). In such cases, EU financial support or compensation schemes may promote collaboration and reduce disparities (European Environment Agency, 2024; Visconti et al., 2024). For environmentally redundant areas, inclusion should be based on additional ecological or local criteria, in line with systematic conservation planning (Margules & Pressey, 2000; Moilanen et al., 2009). While some redundancy can be beneficial, it should be carefully calibrated to ensure efficient use of limited resources.

Even if all core areas and a complementary set of conditional value areas were designated for strict protection, the EU-wide 10% target would remain unattainable. This is even more evident at the level of individual countries, especially in Central and Western Europe, where strict protection coverage is low and roadless areas are virtually absent. This underscores the need to expand the strict protection network beyond roadless areas, even if it entails increased conflict possibilities within productive land. While feasibility considerations are valid, protecting some areas with high opportunity costs remains essential for achieving biodiversity goals (Kloibhofer et al., 2025; Visconti et al., 2024). Degraded landscapes on previously productive areas, especially where natural succession or land abandonment is underway, could be good candidates for strict protection, following restoration or rewilding initiatives to reinstate their roadless character (Ibisch et al., 2016; Perino et al., 2019).

However, such interventions must be context sensitive. In fire-prone areas, for example, a degree of road access may be necessary for safety and management (Johnston et al., 2021), even though roads can increase fire ignition risk, particularly in regions where wildfires are largely human-induced (Kati et al., 2023). Ultimately, a balanced strategy that prioritize roadless areas while considering the inclusion of more infrastructurally developed regions to expand the network of strictly protected areas will be key to meeting conservation targets in the EU.

While our study represents a major advance in the approach for roadless area mapping and prioritization to meet international conservation targets, several limitations must be acknowledged. The reliance of the methodological framework on OSM data, despite its high coverage and frequent updates, may result in regional inconsistencies due to varying mapping completeness and accuracy in the tagging of linear features (Barrington-Leigh & Millard-Ball, 2017). However, as data from European countries are virtually complete, our results can be considered sound. Instead, efforts to apply this framework elsewhere should carefully evaluate its transferability, based on the local availability and quality of OSM data (Hughes, 2017), which could, if necessary, be supplemented with alternative sources. Furthermore, applying a uniform 1 km buffer to define road-effect zones, does not account for taxon- and habitat-specific responses, road types, or terrain features (Ibisch et al., 2016; Wu et al., 2017). Future work should improve precision by developing adaptive buffering models that incorporate these context-specific factors. Lastly, while our prioritization is grounded in environmental representativeness, effective conservation planning must also consider other key biodiversity components such as species-level data, particularly for threatened, endemic, and functionally important taxa, as well as ecosystem and habitat typologies that capture the structural and compositional diversity of natural systems (Margules & Pressey, 2000; Watson et al., 2011). In addition, socioeconomic dimensions, including land tenure security, cultural values, governance capacity, and stakeholder acceptance, are critical for ensuring the feasibility, equity, and long-term sustainability of conservation outcomes (Knight et al., 2006; Redford et al., 2013).

## Conclusions

Our results highlight the pivotal role that remaining roadless areas could play in achieving the EU target of 10% strict protection, particularly by improving the environmental representativeness of the current network. While their overall extent is limited and unevenly distributed across countries, a substantial proportion of roadless areas remains unprotected and could contribute to addressing existing conservation gaps. However, prioritization efforts must account for trade-offs between ecological value, policy feasibility, and spatial equity. Expanding the strict protection network will therefore require a balanced strategy that integrates core and conditional value areas, restoration opportunities, and coordinated action across governance levels to ensure effective and inclusive implementation of conservation targets.

## Data availability statement

An interactive version of the maps presented in this work is available at: https://ee-riccardotestolin-367410.projects.earthengine.app/view/roadless-areas-in-the-european-union. The code and data necessary to replicate the analyses presented in this study, including all preprocessing steps and model outputs, will be made publicly available upon article acceptance. This will include the derived geospatial layers identifying roadless and wilderness areas, environmental space grids, and conservation value classifications.

## Acknowledgements

Project funded under the National Recovery and Resilience Plan (NRRP), Mission 4 Component 2 Investment 1.4 - Call for tender No. 3138 of 16 December 2021, rectified by Decree n.3175 of 18 December 2021 of Italian Ministry of University and Research funded by the European Union – NextGenerationEU; Award Number: Project code CN_00000033, Concession Decree No. 1034 of 17 June 2022 adopted by the Italian Ministry of University and Research, CUP J33C22001190001, Project title “National Biodiversity Future Center - NBFC”. F.M.S. was funded under the Horizon Europe project “FORbEST” (Grant No 10118178). The authors would like to thank Prof. Carlo Rondinini (Sapienza University of Rome) for his insightful comments on the first draft of the manuscript.

