## Supplementary Information for "Leveraging roadless areas to close strict protection gaps in the EU"

**Supplementary information to "Contribution of roadless areas to the strict protection network in the European Union"**

**Table S1.** OSM tags used to identify linear features for the extraction of roadless and wilderness areas. Descriptions are taken from wiki.openstreetmap.org and Frederik Ramm (2022).

| OSM tag | Description | Target |
|---|---|---|
| **Highways** | | |
| highway = motorway | Motorway/freeway | roadless & wilderness |
| highway = trunk | Important roads, typically divided | roadless & wilderness |
| highway = primary | Primary roads, typically national | roadless & wilderness |
| highway = secondary | Secondary roads, typically regional | roadless & wilderness |
| highway = tertiary | Tertiary roads, typically local | roadless & wilderness |
| highway = unclassified | Smaller local roads | roadless & wilderness |
| highway = motorway_link | Roads that connect from one road to another of the same of lower category | roadless & wilderness |
| highway = trunk_link | | roadless & wilderness |
| highway = primary_link | | roadless & wilderness |
| highway = secondary_link | | roadless & wilderness |
| highway = tertiary_link | | roadless & wilderness |
| highway = residential | Roads in residential areas | roadless & wilderness |
| highway = living_street | Streets where pedestrians have priority | roadless & wilderness |
| highway = pedestrian | Pedestrian only streets | roadless & wilderness |
| highway = raceway | A racetrack for motorised racing | roadless & wilderness |
| highway = busway | Dedicated roads for bus | roadless & wilderness |
| highway = bus_guideway | Busway that is side guided "rails like" | roadless & wilderness |
| highway = service | Service roads for access to facilities | roadless & wilderness |
| highway = escape | Emergency ramp | roadless & wilderness |
| highway = cycleway | Paths for cycling | roadless & wilderness |
| highway = construction | Any type of highway under construction | roadless & wilderness |
| highway = road | Road with an unknown classification | wilderness |
| highway = track | For agricultural use, in forests, etc. | wilderness |
| highway = bridleway | Paths for horse riding | wilderness |
| tracktype = grade1 | Tracks can be assigned a track type from 1 (asphalt or heavily compacted) to 5 (hardly visible). A detailed description is here: https://wiki.openstreetmap.org/wiki/Key:tracktype | roadless & wilderness |
| tracktype = grade2 | | roadless & wilderness |
| tracktype = grade3 | | roadless & wilderness |
| tracktype = grade4 | | roadless & wilderness |
| tracktype = grade5 | | roadless & wilderness |
| **Railways** | | |
| railway = rail | Regular railway tracks | roadless & wilderness |
| railway = light_rail | Light railway tracks, often commuter railways | roadless & wilderness |
| railway = tram | Tram tracks (may be incident with roads) | roadless & wilderness |
| railway = monorail | A monorail track | roadless & wilderness |
| railway = narrow_gauge | A narrow-gauge railway track | roadless & wilderness |
| railway = funicular | A funicular, or cable railway | roadless & wilderness |
| railway = rack | A rack railway | roadless & wilderness |
| railway = construction | Any type of railway under construction | roadless & wilderness |
| **Aerialways** | | |
| aerialway = cable_car | A cabin cable car run | wilderness |
| aerialway = gondola | An aerial way with circling cabins | wilderness |
| aerialway = chair_lift | An open chairlift run | wilderness |
| aerialway = drag_lift | An overhead towline for skiers | wilderness |
| aerialway = goods | An aerial way for the transport of goods | wilderness |
| aerialway = t-bar | Other types of lift | wilderness |
| aerialway = j-bar | | wilderness |

| | | |
|---|---|---|
| aerialway = platter | | wilderness |
| aerialway = rope-tow | | wilderness |
| aerialway = magic_carpet | | wilderness |
| aerialway = zipline | | wilderness |
| aerialway = mixed_lift | | wilderness |
| **Waterways** | | |
| waterway = canal | An artificial waterway | wilderness |
| waterway = pressurised | A pressurised artificial conduit | wilderness |
| waterway = drain | A small drainage ditch or similar structure | wilderness |
| waterway = fairway | Navigable route in a body of water | wilderness |
| waterway = fish_pass | A way for fish to circumvent a water barrier | wilderness |
| waterway = canoe_pass | A passage made to cross a dam/weir | wilderness |
| power = cable | An underground or submarine power cable | wilderness |
| power = line | A regular power line | wilderness |
| power = minor_line | A smaller power line | wilderness |
| man_made = pipeline | Major pipelines carrying liquids, gases, etc. | wilderness |
| man_made = goods_conveyor | Permanent conveyor system | wilderness |

**Table S2.** Extent, percentage of country surface (% country) and share (% total) of roadless and wilderness areas for each EU country and the EU27 as a whole.

| | Roadless | | | Wilderness | | |
|---|---|---|---|---|---|---|
| **Country** | **Extent (km$^2$)** | **% country** | **% total** | **Extent (km$^2$)** | **% country** | **% total** |
| Austria | 8,967 | 10.68 | 1.88 | 5,220 | 6.22 | 2.69 |
| Belgium | 38 | 0.12 | 0.01 | 0 | 0 | 0 |
| Bulgaria | 18,788 | 16.84 | 3.94 | 4,108 | 3.68 | 2.12 |
| Croatia | 5,980 | 10.48 | 1.25 | 1,336 | 2.34 | 0.69 |
| Cyprus | 452 | 5.00 | 0.09 | 24 | 0.27 | 0.01 |
| Czechia | 147 | 0.19 | 0.03 | 6 | 0.01 | ~0 |
| Denmark | 787 | 1.77 | 0.17 | 508 | 1.14 | 0.26 |
| Estonia | 3,907 | 8.59 | 0.82 | 2,042 | 4.49 | 1.05 |
| Finland | 82,427 | 24.35 | 17.3 | 42,054 | 12.43 | 21.71 |
| France | 18,011 | 3.28 | 3.78 | 8,167 | 1.49 | 4.22 |
| Germany | 2,392 | 0.67 | 0.5 | 350 | 0.10 | 0.18 |
| Greece | 19,616 | 14.80 | 4.12 | 3,484 | 2.63 | 1.8 |
| Hungary | 2,883 | 3.10 | 0.6 | 41 | 0.04 | 0.02 |
| Ireland | 2,864 | 4.08 | 0.6 | 1,470 | 2.09 | 0.76 |
| Italy | 22,666 | 7.54 | 4.76 | 10,550 | 3.51 | 5.45 |
| Latvia | 2,399 | 3.71 | 0.5 | 1,354 | 2.09 | 0.7 |
| Lituania | 1,572 | 2.42 | 0.33 | 247 | 0.38 | 0.13 |
| Luxembourg | 0 | 0 | 0 | 0 | 0 | 0 |
| Malta | 0 | 0 | 0 | 0 | 0 | 0 |
| Netherlands | 160 | 0.42 | 0.03 | 44 | 0.12 | 0.02 |
| Poland | 7,226 | 2.31 | 1.52 | 400 | 0.13 | 0.21 |
| Portugal | 11,236 | 12.23 | 2.36 | 531 | 0.58 | 0.27 |
| Romania | 44,071 | 18.49 | 9.25 | 12,945 | 5.43 | 6.68 |
| Slovakia | 3,591 | 7.31 | 0.75 | 290 | 0.59 | 0.15 |
| Slovenia | 669 | 3.35 | 0.14 | 358 | 1.79 | 0.18 |
| Spain | 68,866 | 13.61 | 14.45 | 13,749 | 2.72 | 7.1 |
| Sweden | 146,868 | 32.63 | 30.82 | 84,467 | 18.77 | 43.6 |
| **EU27** | **476,583** | **11.51** | **100** | **193,745** | **4.68** | **100** |

**Table S3.** Number (n) and percentage (%) of roadless and wilderness fragments belonging to different size classes for each EU country and the EU27 as a whole. Fragments <1 km$^2$ are the results of the separation of cross-country roadless and wilderness patches.

| | | Roadless | | Wilderness | |
|---|---|---|---|---|---|
| **Country** | **Size class** | **n** | **Area (km$^2$)** | **n** | **Area (km$^2$)** |
| Austria | all | 288 (100%) | 8,968 (100%) | 238 (100%) | 5,221 (100%) |
| | >1 km$^2$ | 279 (96.9%) | 8,965 (100%) | 231 (97.1%) | 5,219 (100%) |
| | >5 km$^2$ | 136 (47.2%) | 8,637 (96%) | 115 (48.3%) | 4,923 (94%) |
| | >10 km$^2$ | 86 (29.9%) | 8,291 (92%) | 75 (31.5%) | 4,625 (89%) |
| | >50 km$^2$ | 28 (9.7%) | 7,055 (79%) | 21 (8.8%) | 3,568 (68%) |
| | >100 km$^2$ | 20 (6.9%) | 6,504 (73%) | 10 (4.2%) | 2,724 (52%) |
| | >1000 km$^2$ | 1 (0.3%) | 1,494 (17%) | 0 | - |
| Belgium | all | 17 (100%) | 38 (100%) | 0 | - |
| | >1 km$^2$ | 16 (94.1%) | 38 (100%) | 0 | - |
| | >5 km$^2$ | 0 | - | 0 | - |
| | >10 km$^2$ | 0 | - | 0 | - |
| | >50 km$^2$ | 0 | - | 0 | - |
| | >100 km$^2$ | 0 | - | 0 | - |
| | >1000 km$^2$ | 0 | - | 0 | - |
| Bulgaria | all | 627 (100%) | 18,788 (100%) | 476 (100%) | 4,108 (100%) |
| | >1 km$^2$ | 624 (99.5%) | 18,788 (100%) | 473 (99.4%) | 4,107 (100%) |
| | >5 km$^2$ | 357 (56.9%) | 18,164 (97%) | 168 (35.3%) | 3,366 (82%) |
| | >10 km$^2$ | 255 (40.7%) | 17,444 (93%) | 85 (17.9%) | 2,775 (68%) |
| | >50 km$^2$ | 81 (12.9%) | 13,268 (71%) | 10 (2.1%) | 1,326 (32%) |
| | >100 km$^2$ | 51 (8.1%) | 11,172 (59%) | 5 (1.1%) | 977 (24%) |
| | >1000 km$^2$ | 0 | - | 0 | - |
| Croatia | all | 592 (100%) | 5,982 (100%) | 190 (100%) | 1,337 (100%) |
| | >1 km$^2$ | 591 (99.8%) | 5,981 (100%) | 190 (100%) | 1,337 (100%) |
| | >5 km$^2$ | 229 (38.7%) | 5,105 (85%) | 64 (33.7%) | 1,057 (79%) |
| | >10 km$^2$ | 132 (22.3%) | 4,428 (74%) | 28 (14.7%) | 800 (60%) |
| | >50 km$^2$ | 27 (4.6%) | 2,270 (38%) | 3 (1.6%) | 325 (24%) |
| | >100 km$^2$ | 4 (0.7%) | 716 (12%) | 1 (0.5%) | 200 (15%) |
| | >1000 km$^2$ | 0 | - | 0 | - |
| Cyprus | all | 69 (100%) | 452 (100%) | 5 (100%) | 24 (100%) |
| | >1 km$^2$ | 69 (100%) | 452 (100%) | 5 (100%) | 24 (100%) |
| | >5 km$^2$ | 19 (27.5%) | 325 (72%) | 1 (20%) | 15 (63%) |
| | >10 km$^2$ | 8 (11.6%) | 241 (53%) | 1 (20%) | 15 (63%) |
| | >50 km$^2$ | 1 (1.4%) | 66 (15%) | 0 | - |
| | >100 km$^2$ | 0 | - | 0 | - |
| | >1000 km$^2$ | 0 | - | 0 | - |
| Czechia | all | 58 (100%) | 146 (100%) | 3 (100%) | 5 (94%) |
| | >1 km$^2$ | 51 (87.9%) | 145 (99%) | 1 (33.3%) | 5 (94%) |
| | >5 km$^2$ | 9 (15.5%) | 71 (49%) | 0 | - |
| | >10 km$^2$ | 2 (3.4%) | 25 (17%) | 0 | - |
| | >50 km$^2$ | 0 | - | 0 | - |
| | >100 km$^2$ | 0 | - | 0 | - |
| | >1000 km$^2$ | 0 | - | 0 | - |
| Denmark | all | 87 (100%) | 787 (100%) | 54 (100%) | 508 (100%) |
| | >1 km$^2$ | 87 (100%) | 787 (100%) | 54 (100%) | 508 (100%) |
| | >5 km$^2$ | 44 (50.6%) | 688 (87%) | 34 (63%) | 461 (91%) |
| | >10 km$^2$ | 23 (26.4%) | 548 (70%) | 17 (31.5%) | 350 (69%) |

| | | | | | |
|---|---|---|---|---|---|
| | >50 km$^2$ | 2 (2.3%) | 196 (25%) | 2 (3.7%) | 135 (27%) |
| | >100 km$^2$ | 1 (1.1%) | 117 (15%) | 0 | - |
| | >1000 km$^2$ | 0 | - | 0 | - |
| Estonia | all | 415 (100%) | 3,908 (100%) | 210 (100%) | 2,042 (100%) |
| | >1 km$^2$ | 413 (99.5%) | 3,907 (100%) | 208 (99%) | 2,041 (100%) |
| | >5 km$^2$ | 155 (37.3%) | 3,288 (84%) | 72 (34.3%) | 1,722 (84%) |
| | >10 km$^2$ | 75 (18.1%) | 2,723 (70%) | 41 (19.5%) | 1,497 (73%) |
| | >50 km$^2$ | 15 (3.6%) | 1,526 (39%) | 10 (4.8%) | 859 (42%) |
| | >100 km$^2$ | 6 (1.4%) | 882 (23%) | 2 (1%) | 249 (12%) |
| | >1000 km$^2$ | 0 | - | 0 | - |
| Finland | all | 3,120 (100%) | 82,428 (100%) | 1,851 (100%) | 42,054 (100%) |
| | >1 km$^2$ | 3,118 (99.9%) | 82,427 (100%) | 1,849 (99.9%) | 42,054 (100%) |
| | >5 km$^2$ | 1,397 (44.8%) | 78,460 (95%) | 691 (37.3%) | 39,407 (94%) |
| | >10 km$^2$ | 875 (28%) | 74,777 (91%) | 398 (21.5%) | 37,323 (89%) |
| | >50 km$^2$ | 246 (7.9%) | 61,382 (74%) | 73 (3.9%) | 30,528 (73%) |
| | >100 km$^2$ | 102 (3.3%) | 51,368 (62%) | 44 (2.4%) | 28,402 (68%) |
| | >1000 km$^2$ | 8 (0.3%) | 29,287 (36%) | 10 (0.5%) | 19,209 (46%) |
| France | all | 1,291 (100%) | 18,011 (100%) | 279 (100%) | 8,167 (100%) |
| | >1 km$^2$ | 1,285 (99.5%) | 18,008 (100%) | 276 (98.9%) | 8,166 (100%) |
| | >5 km$^2$ | 395 (30.6%) | 15,963 (89%) | 127 (45.5%) | 7,816 (96%) |
| | >10 km$^2$ | 210 (16.3%) | 14,671 (81%) | 91 (32.6%) | 7,554 (92%) |
| | >50 km$^2$ | 52 (4%) | 11,424 (63%) | 39 (14%) | 6,318 (77%) |
| | >100 km$^2$ | 33 (2.6%) | 10,000 (56%) | 23 (8.2%) | 5,175 (63%) |
| | >1000 km$^2$ | 1 (0.1%) | 1,293 (7%) | 0 | - |
| Germany | all | 444 (100%) | 2,393 (100%) | 50 (100%) | 351 (100%) |
| | >1 km$^2$ | 441 (99.3%) | 2,392 (100%) | 48 (96%) | 349 (100%) |
| | >5 km$^2$ | 114 (25.7%) | 1,652 (69%) | 18 (36%) | 282 (80%) |
| | >10 km$^2$ | 48 (10.8%) | 1,196 (50%) | 7 (14%) | 200 (57%) |
| | >50 km$^2$ | 7 (1.6%) | 457 (19%) | 1 (2%) | 92 (26%) |
| | >100 km$^2$ | 1 (0.2%) | 109 (5%) | 0 | - |
| | >1000 km$^2$ | 0 | - | 0 | - |
| Greece | all | 1,406 (100%) | 19,616 (100%) | 629 (100%) | 3,484 (100%) |
| | >1 km$^2$ | 1,406 (100%) | 19,616 (100%) | 625 (99.4%) | 3,483 (100%) |
| | >5 km$^2$ | 626 (44.5%) | 17,680 (90%) | 158 (25.1%) | 2,373 (68%) |
| | >10 km$^2$ | 385 (27.4%) | 15,982 (81%) | 63 (10%) | 1,708 (49%) |
| | >50 km$^2$ | 85 (6%) | 9,776 (50%) | 6 (1%) | 658 (19%) |
| | >100 km$^2$ | 32 (2.3%) | 6,118 (31%) | 2 (0.3%) | 373 (11%) |
| | >1000 km$^2$ | 0 | - | 0 | - |
| Hungary | all | 359 (100%) | 2,883 (100%) | 5 (100%) | 41 (100%) |
| | >1 km$^2$ | 354 (98.6%) | 2,882 (100%) | 5 (100%) | 41 (100%) |
| | >5 km$^2$ | 148 (41.2%) | 2,416 (84%) | 1 (20%) | 34 (83%) |
| | >10 km$^2$ | 81 (22.6%) | 1,944 (67%) | 1 (20%) | 34 (83%) |
| | >50 km$^2$ | 7 (2%) | 494 (17%) | 0 | - |
| | >100 km$^2$ | 1 (0.3%) | 117 (4%) | 0 | - |
| | >1000 km$^2$ | 0 | - | 0 | - |
| Ireland | all | 211 (100%) | 2,864 (100%) | 142 (100%) | 1,470 (100%) |
| | >1 km$^2$ | 211 (100%) | 2,864 (100%) | 142 (100%) | 1,470 (100%) |
| | >5 km$^2$ | 101 (47.9%) | 2,610 (91%) | 70 (49.3%) | 1,284 (87%) |
| | >10 km$^2$ | 68 (32.2%) | 2,369 (83%) | 46 (32.4%) | 1,105 (75%) |
| | >50 km$^2$ | 12 (5.7%) | 1,011 (35%) | 4 (2.8%) | 272 (19%) |
| | >100 km$^2$ | 2 (0.9%) | 331 (12%) | 0 | - |

| | | | | | |
|---|---|---|---|---|---|
| | >1000 km$^2$ | 0 | - | 0 | - |
| Italy | all | 1,278 (100%) | 22,667 (100%) | 561 (100%) | 10,551 (100%) |
| | >1 km$^2$ | 1,277 (99.9%) | 22,667 (100%) | 556 (99.1%) | 10,549 (100%) |
| | >5 km$^2$ | 522 (40.8%) | 20,874 (92%) | 244 (43.5%) | 9,798 (93%) |
| | >10 km$^2$ | 323 (25.3%) | 19,456 (86%) | 162 (28.9%) | 9,231 (87%) |
| | >50 km$^2$ | 83 (6.5%) | 14,576 (64%) | 46 (8.2%) | 6,627 (63%) |
| | >100 km$^2$ | 42 (3.3%) | 11,702 (52%) | 20 (3.6%) | 4,917 (47%) |
| | >1000 km$^2$ | 1 (0.1%) | 1,516 (7%) | 0 | - |
| Latvia | all | 384 (100%) | 2,399 (100%) | 201 (100%) | 1,355 (100%) |
| | >1 km$^2$ | 378 (98.4%) | 2,397 (100%) | 200 (99.5%) | 1,354 (100%) |
| | >5 km$^2$ | 100 (26%) | 1,775 (74%) | 61 (30.3%) | 1,067 (79%) |
| | >10 km$^2$ | 51 (13.3%) | 1,455 (61%) | 28 (13.9%) | 829 (61%) |
| | >50 km$^2$ | 7 (1.8%) | 661 (28%) | 3 (1.5%) | 375 (28%) |
| | >100 km$^2$ | 2 (0.5%) | 361 (15%) | 2 (1%) | 320 (24%) |
| | >1000 km$^2$ | 0 | - | 0 | - |
| Lithuania | all | 331 (100%) | 1,572 (100%) | 52 (100%) | 247 (100%) |
| | >1 km$^2$ | 328 (99.1%) | 1,571 (100%) | 51 (98.1%) | 246 (100%) |
| | >5 km$^2$ | 80 (24.2%) | 1,040 (66%) | 8 (15.4%) | 161 (65%) |
| | >10 km$^2$ | 30 (9.1%) | 703 (45%) | 4 (7.7%) | 132 (54%) |
| | >50 km$^2$ | 3 (0.9%) | 220 (14%) | 0 | - |
| | >100 km$^2$ | 0 | - | 0 | - |
| | >1000 km$^2$ | 0 | - | 0 | - |
| Luxembourg | all | 0 | - | 0 | - |
| | >1 km$^2$ | 0 | - | 0 | - |
| | >5 km$^2$ | 0 | - | 0 | - |
| | >10 km$^2$ | 0 | - | 0 | - |
| | >50 km$^2$ | 0 | - | 0 | - |
| | >100 km$^2$ | 0 | - | 0 | - |
| | >1000 km$^2$ | 0 | - | 0 | - |
| Malta | all | 0 | - | 0 | - |
| | >1 km$^2$ | 0 | - | 0 | - |
| | >5 km$^2$ | 0 | - | 0 | - |
| | >10 km$^2$ | 0 | - | 0 | - |
| | >50 km$^2$ | 0 | - | 0 | - |
| | >100 km$^2$ | 0 | - | 0 | - |
| | >1000 km$^2$ | 0 | - | 0 | - |
| Netherlands | all | 34 (100%) | 159 (100%) | 15 (100%) | 44 (101%) |
| | >1 km$^2$ | 34 (100%) | 159 (100%) | 15 (100%) | 44 (101%) |
| | >5 km$^2$ | 10 (29.4%) | 103 (65%) | 1 (6.7%) | 6 (14%) |
| | >10 km$^2$ | 4 (11.8%) | 66 (42%) | 0 | - |
| | >50 km$^2$ | 0 | - | 0 | - |
| | >100 km$^2$ | 0 | - | 0 | - |
| | >1000 km$^2$ | 0 | - | 0 | - |
| Poland | all | 1,441 (100%) | 7,225 (100%) | 66 (100%) | 400 (100%) |
| | >1 km$^2$ | 1,434 (99.5%) | 7,223 (100%) | 65 (98.5%) | 399 (100%) |
| | >5 km$^2$ | 356 (24.7%) | 4,793 (66%) | 15 (22.7%) | 311 (78%) |
| | >10 km$^2$ | 148 (10.3%) | 3,385 (47%) | 9 (13.6%) | 268 (67%) |
| | >50 km$^2$ | 10 (0.7%) | 890 (12%) | 0 | - |
| | >100 km$^2$ | 3 (0.2%) | 497 (7%) | 0 | - |
| | >1000 km$^2$ | 0 | - | 0 | - |
| Portugal | all | 957 (100%) | 11,237 (100%) | 141 (100%) | 532 (100%) |

| | | | | | |
|---|---|---|---|---|---|
| | >1 km² | 954 (99.7%) | 11,236 (100%) | 136 (96.5%) | 529 (99%) |
| | >5 km² | 446 (46.6%) | 9,997 (89%) | 28 (19.9%) | 306 (58%) |
| | >10 km² | 268 (28%) | 8,745 (78%) | 9 (6.4%) | 174 (33%) |
| | >50 km² | 50 (5.2%) | 4,355 (39%) | 1 (0.7%) | 72 (14%) |
| | >100 km² | 12 (1.3%) | 1,901 (17%) | 0 | - |
| | >1000 km² | 0 | - | 0 | - |
| Romania | all | 1,372 (100%) | 44,071 (100%) | 1,278 (100%) | 12,945 (100%) |
| | >1 km² | 1,371 (99.9%) | 44,070 (100%) | 1,278 (100%) | 12,945 (100%) |
| | >5 km² | 755 (55%) | 42,545 (97%) | 514 (40.2%) | 11,120 (86%) |
| | >10 km² | 519 (37.8%) | 40,892 (93%) | 272 (21.3%) | 9,392 (73%) |
| | >50 km² | 152 (11.1%) | 32,889 (75%) | 42 (3.3%) | 4,734 (37%) |
| | >100 km² | 89 (6.5%) | 28,632 (65%) | 15 (1.2%) | 2,834 (22%) |
| | >1000 km² | 4 (0.3%) | 5,528 (13%) | 0 | - |
| Slovakia | all | 520 (100%) | 3,591 (100%) | 27 (100%) | 290 (100%) |
| | >1 km² | 514 (98.8%) | 3,588 (100%) | 26 (96.3%) | 289 (100%) |
| | >5 km² | 184 (35.4%) | 2,816 (78%) | 6 (22.2%) | 244 (84%) |
| | >10 km² | 89 (17.1%) | 2,160 (60%) | 4 (14.8%) | 229 (79%) |
| | >50 km² | 6 (1.2%) | 618 (17%) | 2 (7.4%) | 199 (69%) |
| | >100 km² | 1 (0.2%) | 333 (9%) | 1 (3.7%) | 119 (41%) |
| | >1000 km² | 0 | - | 0 | - |
| Slovenia | all | 60 (100%) | 670 (100%) | 18 (100%) | 358 (100%) |
| | >1 km² | 56 (93.3%) | 668 (100%) | 17 (94.4%) | 358 (100%) |
| | >5 km² | 15 (25%) | 575 (86%) | 8 (44.4%) | 333 (93%) |
| | >10 km² | 8 (13.3%) | 534 (80%) | 5 (27.8%) | 314 (88%) |
| | >50 km² | 3 (5%) | 419 (63%) | 2 (11.1%) | 256 (72%) |
| | >100 km² | 1 (1.7%) | 284 (42%) | 1 (5.6%) | 199 (56%) |
| | >1000 km² | 0 | - | 0 | - |
| Spain | all | 3,692 (100%) | 68,867 (100%) | 1,608 (100%) | 13,750 (100%) |
| | >1 km² | 3,680 (99.7%) | 68,862 (100%) | 1,603 (99.7%) | 13,747 (100%) |
| | >5 km² | 1,931 (52.3%) | 64,490 (94%) | 592 (36.8%) | 11,388 (83%) |
| | >10 km² | 1,308 (35.4%) | 60,082 (87%) | 296 (18.4%) | 9,274 (67%) |
| | >50 km² | 318 (8.6%) | 38,294 (56%) | 35 (2.2%) | 4,097 (30%) |
| | >100 km² | 138 (3.7%) | 25,727 (37%) | 12 (0.7%) | 2,607 (19%) |
| | >1000 km² | 1 | 1538 (2%) | 0 | - |
| Sweden | all | 3,894 (100%) | 1468,69 (100%) | 1,986 (100%) | 84,468 (100%) |
| | >1 km² | 3,891 (99.9%) | 1468,69 (100%) | 1,984 (99.9%) | 84,466 (100%) |
| | >5 km² | 2,040 (52.4%) | 142,458 (97%) | 692 (34.8%) | 81,505 (96%) |
| | >10 km² | 1,369 (35.2%) | 137,678 (94%) | 369 (18.6%) | 79,178 (94%) |
| | >50 km² | 357 (9.2%) | 114,944 (78%) | 95 (4.8%) | 73,688 (87%) |
| | >100 km² | 149 (3.8%) | 100,354 (68%) | 61 (3.1%) | 71,317 (84%) |
| | >1000 km² | 14 (0.4%) | 68,044 (46%) | 11 (0.6%) | 56,167 (66%) |
| **EU27** | **>1 km²** | **22,701 (100%)** | **476,589 (100%)** | **9,967 (100%)** | **193,748 (100%)** |
| | **>5 km²** | **10,089 (44.4%)** | **446,749 (94%)** | **3,658 (36.7%)** | **179,108 (92%)** |
| | **>10 km²** | **6,302 (27.8%)** | **420,136 (88%)** | **1,984 (19.9%)** | **167,182 (86%)** |
| | **>50 km²** | **1,531 (6.7%)** | **318,101 (67%)** | **381 (3.8%)** | **134,627 (69%)** |
| | **>100 km²** | **678 (3%)** | **259,144 (54%)** | **199 (2%)** | **121,944 (63%)** |
| | **>1000 km²** | **31 (0.1%)** | **111,368 (23%)** | **22 (0.2%)** | **76,459 (39%)** |

**Table S4.** Extent and proportion (%) of roadless and wilderness areas under strict protection (IUCN categories I & II). Percentage of country surface under strict protection (including developed areas) and projected increase if wilderness and roadless areas were included.

| | Roadless in strict PA | | Wilderness in strict PA | | % country | | |
|---|---|---|---|---|---|---|---|
| **Country** | **Extent ($km^2$)** | **%** | **Extent ($km^2$)** | **%** | **Strict PA** | **Strict PA + wilderness** | **Strict PA + roadless** |
| Austria | 1,405 | 15.67 | 1,040 | 19.92 | 2.57 | 7.55 | 11.58 |
| Belgium | 0 | 0 | - | - | 0.88 | 0.88 | 1.00 |
| Bulgaria | 1,679 | 8.94 | 1,087 | 26.46 | 2.00 | 4.71 | 17.34 |
| Croatia | 370 | 6.19 | 187 | 14.00 | 3.17 | 5.19 | 13.00 |
| Cyprus | 38 | 8.41 | 2 | 8.33 | 3.28 | 3.52 | 7.86 |
| Czechia | 12 | 8.16 | 1 | 16.67 | 1.16 | 1.17 | 1.33 |
| Denmark | 32 | 4.07 | 5 | 0.98 | 0.20 | 1.33 | 1.89 |
| Estonia | 1,027 | 26.29 | 799 | 39.13 | 3.54 | 6.27 | 9.87 |
| Finland | 28,998 | 35.18 | 25,479 | 60.59 | 9.63 | 14.53 | 25.42 |
| France | 2,275 | 12.63 | 1,791 | 21.93 | 0.81 | 1.97 | 3.67 |
| Germany | 243 | 10.16 | 117 | 33.43 | 0.60 | 0.66 | 1.20 |
| Greece | 484 | 2.47 | 252 | 7.23 | 0.72 | 3.15 | 15.15 |
| Hungary | 280 | 9.71 | 34 | 82.93 | 2.32 | 2.33 | 5.12 |
| Ireland | 373 | 13.02 | 243 | 16.53 | 1.00 | 2.75 | 4.55 |
| Italy | 4,240 | 18.71 | 2,147 | 20.35 | 5.19 | 7.98 | 11.32 |
| Latvia | 497 | 20.72 | 294 | 21.71 | 5.69 | 7.33 | 8.63 |
| Lithuania | 338 | 21.5 | 118 | 47.77 | 2.72 | 2.91 | 4.61 |
| Luxembourg | - | - | - | - | 4.30 | 4.30 | 4.30 |
| Malta | - | - | - | - | 0.93 | 0.93 | 0.93 |
| Netherlands | 57 | 35.62 | 16 | 36.36 | 4.40 | 4.47 | 4.67 |
| Poland | 562 | 7.78 | 208 | 52.00 | 0.64 | 0.70 | 2.77 |
| Portugal | 246 | 2.19 | 124 | 23.35 | 0.87 | 1.31 | 12.83 |
| Romania | 2,104 | 4.77 | 1,006 | 7.77 | 1.34 | 6.34 | 18.94 |
| Slovakia | 515 | 14.34 | 236 | 81.38 | 2.83 | 2.94 | 9.10 |
| Slovenia | 346 | 51.72 | 277 | 77.37 | 4.38 | 4.79 | 6.00 |
| Spain | 3,092 | 4.49 | 1,279 | 9.30 | 1.89 | 4.35 | 14.89 |
| Sweden | 40,913 | 27.86 | 37,491 | 44.39 | 10.64 | 21.08 | 34.19 |
| **EU27** | **90,126** | **18.91** | **74,233** | **38.31** | **3.40** | **6.29** | **12.73** |

**Table S5.** Proportion of roadless (including wilderness) areas in each conservation priority class relative to the total roadless area falling outside the strict protection network. The potential percentage increase in strictly protected area coverage if those areas is reported in bracket. The figures are reported also for those areas that could not be classified due to the lack of environmental information.

| **Country** | **Core priority (%)** | **Conditional priority (%)** | **Low priority (%)** | **Not applicable (%)** |
|---|---|---|---|---|
| Austria | 0.33 (+0.03) | 44.4 (+4) | 55.26 (+4.98) | 0.01 (~+0) |
| Belgium | 13.16 (+0.02) | 86.84 (+0.11) | 0 | 0 |
| Bulgaria | 5.63 (+0.86) | 94.37 (+14.47) | 0 | 0.01 (~+0) |
| Croatia | 3.76 (+0.37) | 95.31 (+9.37) | 0 | 0.93 (+0.09) |
| Cyprus | 0.48 (+0.02) | 98.79 (+4.52) | 0 | 0.73 (+0.03) |
| Czechia | 27.61 (+0.05) | 65.67 (+0.11) | 6.72 (+0.01) | 0 |
| Denmark | 12.04 (+0.2) | 82.54 (+1.4) | 0 | 5.42 (+0.09) |
| Estonia | 28.53 (+1.81) | 69.25 (+4.39) | 1.49 (+0.09) | 0.73 (+0.05) |
| Finland | 4.78 (+0.76) | 42.63 (+6.73) | 52.26 (+8.25) | 0.34 (+0.05) |
| France | 20.35 (+0.58) | 62.3 (+1.78) | 17.06 (+0.49) | 0.29 (+0.01) |
| Germany | 44.7 (+0.27) | 52.56 (+0.32) | 1.86 (+0.01) | 0.88 (+0.01) |
| Greece | 8.37 (+1.21) | 90.42 (+13.05) | 0.06 (+0.01) | 1.14 (+0.17) |
| Hungary | 14.06 (+0.39) | 85.59 (+2.39) | 0.35 (+0.01) | 0 |
| Ireland | 2.81 (+0.1) | 82.1 (+2.91) | 13.65 (+0.48) | 1.45 (+0.05) |
| Italy | 1.43 (+0.09) | 70.56 (+4.32) | 27.85 (+1.71) | 0.16 (+0.01) |
| Latvia | 70.35 (+2.07) | 29.65 (+0.87) | 0 | 0 |
| Lithuania | 38.41 (+0.73) | 60.78 (+1.15) | 0.65 (+0.01) | 0.16 (~+0) |
| Luxembourg | - | - | - | - |
| Malta | - | - | - | - |
| Netherlands | 8.74 (+0.02) | 66.99 (+0.18) | 0 | 24.27 (+0.07) |
| Poland | 57.5 (+1.23) | 39.85 (+0.85) | 2.42 (+0.05) | 0.24 (+0.01) |
| Portugal | 3.01 (+0.36) | 96.57 (+11.55) | 0.27 (+0.03) | 0.15 (+0.02) |
| Romania | 3.12 (+0.55) | 91.35 (+16.08) | 5.49 (+0.97) | 0.05 (+0.01) |
| Slovakia | 1.82 (+0.11) | 82.48 (+5.17) | 15.7 (+0.98) | 0 |
| Slovenia | 2.78 (+0.05) | 47.22 (+0.77) | 50 (+0.81) | 0 |
| Spain | 9.82 (+1.28) | 89.99 (+11.7) | 0.17 (+0.02) | 0.02 (~+0) |
| Sweden | 5.37 (+1.26) | 32.42 (+7.63) | 62.07 (+14.61) | 0.15 (+0.03) |
| **EU27** | **7.94 (+0.74)** | **63.52 (+5.93)** | **28.31 (+2.64)** | **0.23 (+0.02)** |

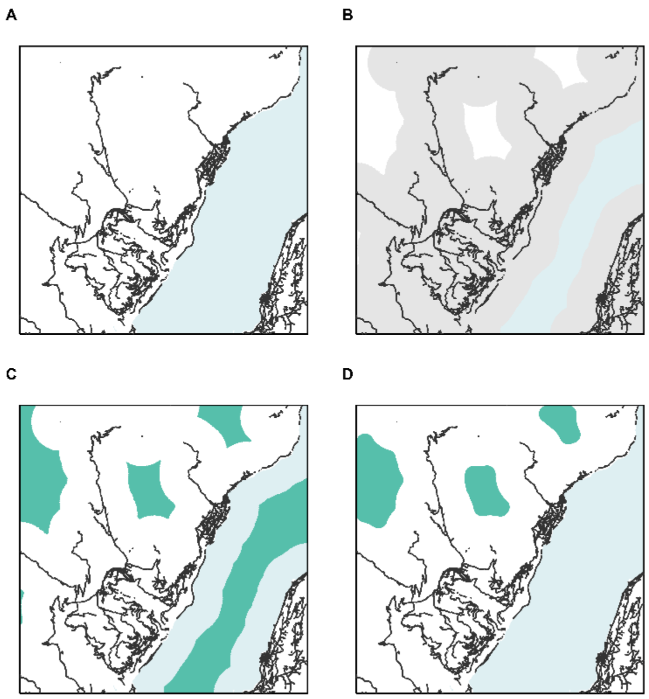


**Figure S1. Workflow for the identification of roadless areas.** A) Road features extraction from OSM. B) 1 km buffer around features. C) Extraction of roadless areas beyond buffer. D) Exclusion of edges and removal of water bodies.

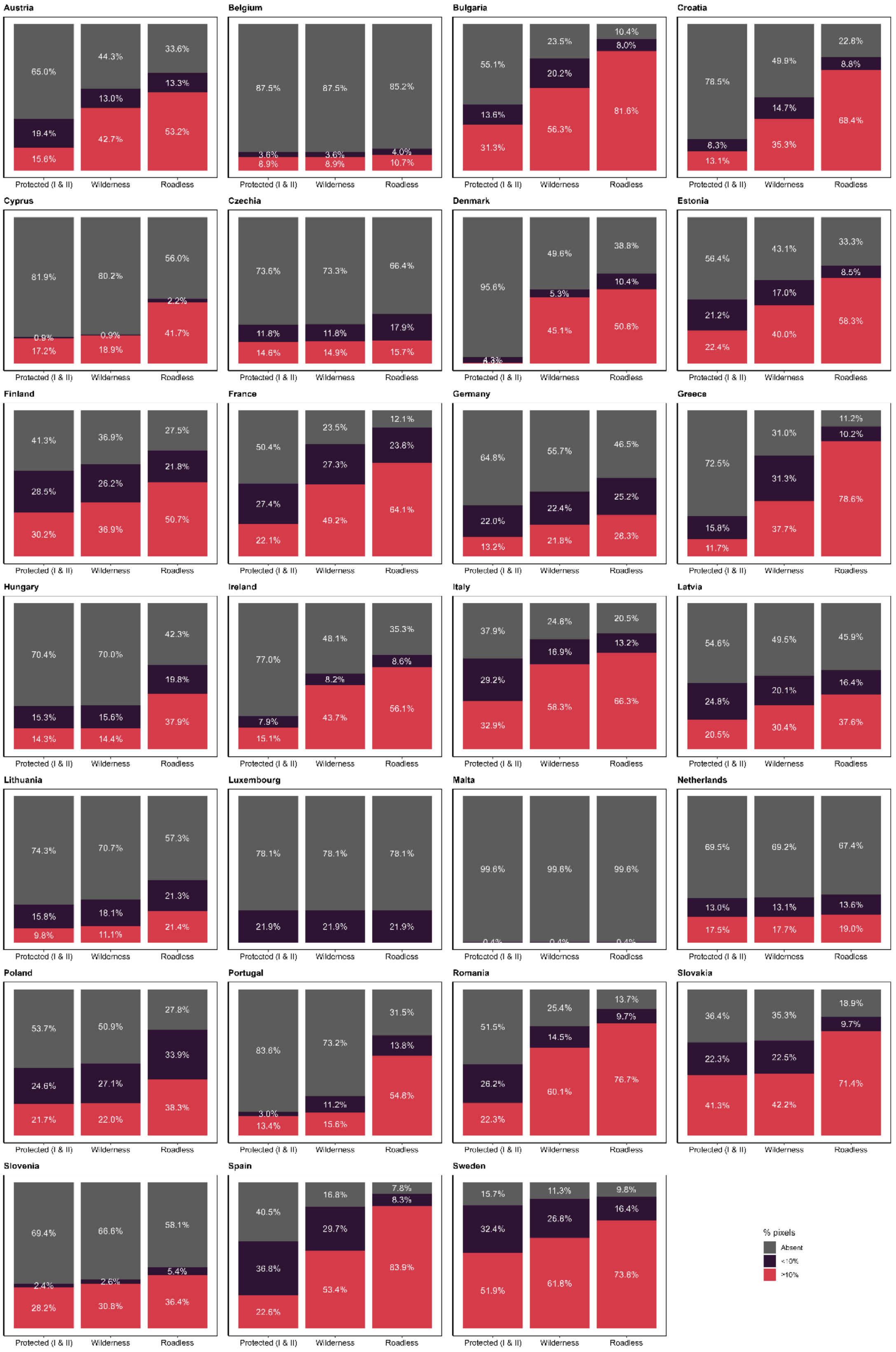


**Figure S2. Representativeness of the environmental space of individual countries.** Proportion of the environmental space (i.e., percentage of grid cells in the principal component plane) of each EU country covered by the current strict protection network and the hypothetical configurations including all wilderness and roadless areas. A distinction between overrepresented and underrepresented cells – when the proportion of pixels included by the network are above and below the 10% threshold, respectively – is provided.
